# Period Doubling Bifurcations in a Forced Cell-Free Genetic Oscillator

**DOI:** 10.1101/2021.12.15.472802

**Authors:** Lukas Aufinger, Johann Brenner, Friedrich C. Simmel

**Affiliations:** Physics Department E14, D-85748 Garching, Germany

## Abstract

Complex non-linear dynamics such as period doubling and chaos have been previously found in computational models of the oscillatory gene networks of biological circadian clocks, but their experimental study is difficult. Here, we present experimental evidence of period doubling in a forced synthetic genetic oscillator operated in a cell-free gene expression system. To this end, an oscillatory negative feedback gene circuit is established in a microfluidic reactor, which allows continuous operation of the system over extended periods of time. We first thoroughly characterize the unperturbed oscillator and find good agreement with a four-species ODE model of the system. Guided by simulations, microfluidics is then used to periodically perturb the system by modulating the concentration of one of the oscillator components with a given amplitude and frequency. When the ratio of the external ‘zeitgeber’ period and the intrinisic period is close to 1, we experimentally find period doubling and quadrupling in the oscillator dynamics, whereas for longer zeitgeber periods, we find stable entrainment. Our theoretical model suggests favorable conditions for which the oscillator can be utilized as an externally synchronized clock, but also demonstrates that related systems could, in principle, display chaotic dynamics.

## I. INTRODUCTION

Across biological kingdoms, organisms including bacteria, fungi, plants, insects, and humans, regulate their day-night cycle with a circadian clock [1, 2]. The ability to measure time is presumed to have concrete evolutionary advantages [2]. In humans, malfunctions of the circadian clock are associated with diseases such as sleep disorders [3], or cancer [4]. At the molecular level, circadian clocks are often comprised of coupled genetic oscillators that are synchronized to external zeitgeber signals [5]. Theoretical studies of various circadian oscillators predict that both forced [6, 7] and freely coupled systems [8, 9] exhibit non-linear phenomena such as splitting, period-doubling, and chaos. While there is experimental evidence for de-synchronization [10] and splitting [11], observation of period-doubling and chaos in circadian clocks have remained elusive due to the experimental challenges associated with long-term observations of such systems [9].

Experimental investigation of period-doubling and chaos in a biological organism would require the accurate measurement of amplitude over many oscillation periods in a potentially fluctuating environment and in the presence of homeostatic regulation processes. One strategy to circumvent these challenges is to study minimal synthetic gene oscillators that can be operated in a controlled and isolated context. Synthetic oscillators have been previously implemented in bacteria [12], mammalian cells [13], and in cell-free batch [14, 15] or continuous reactions [16, 17]. Such systems have been used to study synchronization between communicating cells [18, 19] and among coupled oscillators [20], but also the effects of partitioning [21] and gene expression noise [22]. Transient oscillations have been found close to bifurcations [23].

Engineered gene oscillators can provide molecular rhythms or act as biochemical clocks in other contexts than their circadian counterparts. For instance, the oscillation period of a synthetic oscillator has been used as an accurate measure of bacterial growth rate [22, 24]. Cell-free gene oscillators have been utilized to drive autonomous molecular devices [25], control self-assembly processes [26] or spatio-temporal pattern formation [19]. Previously established synthetic oscillators were operated without periodic synchronization to an external signal, however, and thus provided only an intrinsic measure of time, which lost synchrony with ‘universal time’ after a few periods [27].

Here, we investigate the synchronization of a cell-free genetic oscillator [19, 20] to an external zeitgeber signal using a microfluidic reactor [16] that was previously employed for rapid prototyping of gene circuits [28]. We first verified that the dynamics of the free-running oscillator are well described by a simple model comprised of only four ordinary differential equations (ODEs). We then tested the effects of periodic forcing on the oscillator within the model, and found that the system displays period doubling bifurcations when varying the ratio of the input period to the period of the free oscillator λ = *T_in_/T* in the simulations.

Experimentally, we realized the external forcing by periodically adding either a transcriptional repressor (TetR) or an inducer (aTc), and recording experimental time traces for up to 48 hours. For input periods close to the intrinsic period of the oscillator (λ ≈ 1), we indeed find evidence of period doubling and even quadrupling in the forced system. Larger values of λ result in stable 1-cycles ‘entrained’ on the external zeitgeber. Further analysis, aided by simulations, suggests that with increasing non-linearity in the biochemical feedback loop, similar driven systems could display increasingly complex dynamics, including chaos.

## II. RESULTS

### A. ODE model of the oscillator circuit

As shown in Fig. 1(a), our oscillator circuit consists of two regulatory proteins. Sigma factor *σ*^28^ activates the expression of TetR, which in turn represses the expression of the activator, thereby forming a negative feedback loop. In the experiment, the dynamics of the system is monitored by co-expression of the fluorescent reporter proteins mVenus and mTurquoise2 for the activator and repressor, respectively. To synchronize the oscillator to an external clock signal, the system can be perturbed by either adding purified TetR from the outside, or by inactivating intrinsic TetR via induction with anhydrotetracycline (aTc). Already in this coarse-grained picture the system is constituted of three coupled dynamical variables - activator, inhibitor, and external signal -, which is one of the requirements for a system to exhibit complex non-linear dynamics [29].

**FIG. 1.**
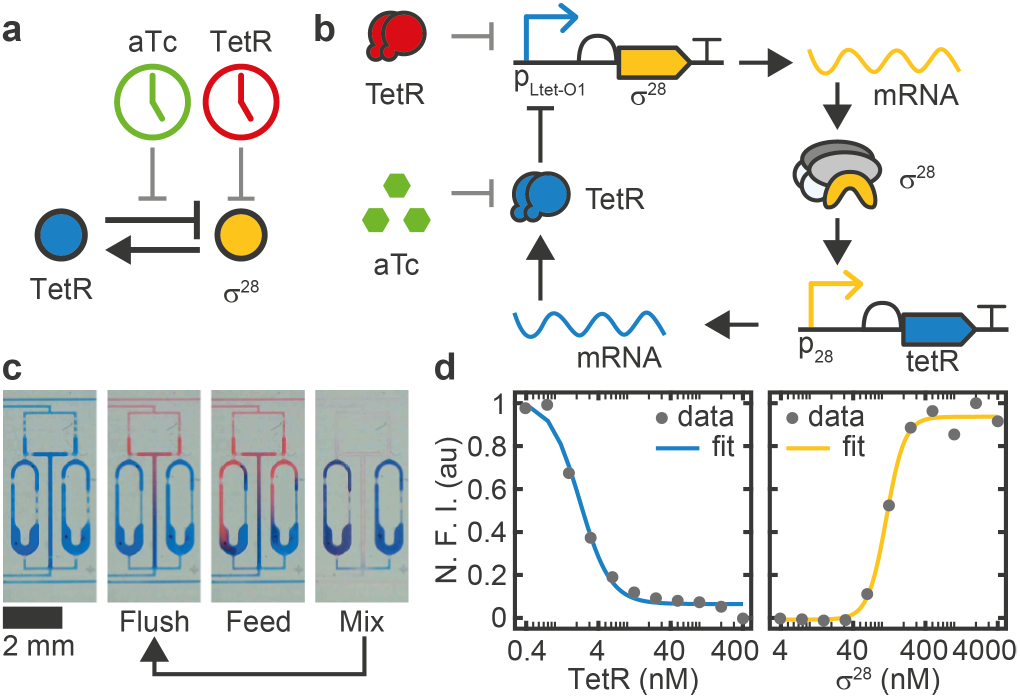
Genetic oscillator circuit and experimental setup. (a) Simple and (b) detailed circuit diagram of the synthetic genetic oscillator consisting of a negative feedback loop where *σ*^28^ acts as activator and TetR acts as repressor. Our four-variable ODE model (Eq. (1)–(4)) considers the concentrations of the two proteins and their respective mRNAs. Protein expression is monitored via co-expression of fluorescent reporters mVenus and mTurquoise2, respectively. We can perturb the system either by inactivating existing TetR by induction with aTc, or by adding purified TetR. (c) A microfluidic ring reactor [16] was used to maintain the reaction out-of-equilibrium by periodically exchanging a fraction of the reactor volume with fresh reagents. By switching between different input reagents, the reactions can be exposed to an arbitrary series of inputs. (d) Transfer functions of the two promoters determined by titrating the regulator protein in bulk. Fitted Hill parameters with 68% confidence intervals are *K_h_* = 2.2 ± 0.2 nM, *n_h_* = 2.1 ± 0.3, *K_a_* = 115 ± 6 nM *n_a_* = 3.4 ± 0.6. N.F.I.: Normalized Fluorescence Intensity.

To properly describe the dynamics of the genetic oscillator, however, it is necessary to explicitly consider the dynamics of the mRNA molecules (Fig. 1(b)), which effectively generates the time delay that is required for sustained oscillations [30]. The dynamics of the free oscillator is then described by the following set of four ordinary differential equations,

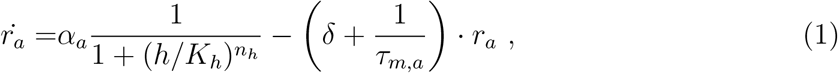

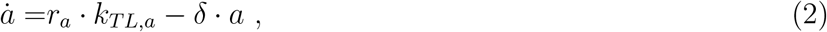

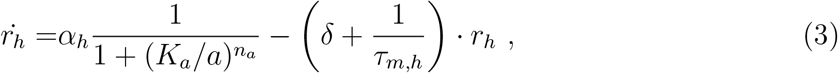

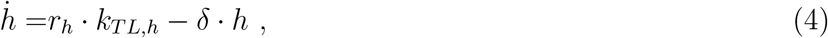

which has been used previously for the analysis of similar systems [15, 19]. The variables *r_a_*, *a*, *r_h_* and *h* denote the concentrations of the activator and inhibitor mRNA and protein species, respectively. To aid the following discussion, we conceptually distinguish between *system parameters* that are essentially fixed properties depending on molecular details, and *control parameters* that can be experimentally tuned relatively easily.

The transcription rates *α_a_* and *α_h_* can be tuned linearly by adjusting the gene template concentrations [16, 31] and will therefore be considered as control parameters. A third control parameter is given by the dilution rate *δ*, which, as shown below, defines the timescale of the system. Experimentally, the reaction is kept out of equilibrium using a semi-continuously operated microfluidic ring reactor [16]. As shown in Fig. 1(c), the reactions are maintained inside ring-like reaction chambers, whose volume is periodically replaced by a fraction *R* (0 < *R* < 1) of fresh reagents, called the ‘refresh ratio’. With a fixed time interval *t_int_* between dilution cycles, the dilution rate can be precisely tuned by varying *R* according to

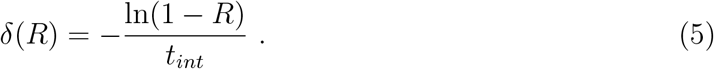

It had been previously shown that a reactor operated in semi-continuous mode can be used to emulate reactions in a continuously operated flow reactor, provided that *t_int_* is sufficiently small compared to the systems timescale [16, 28].

The system parameters are the mRNA lifetimes *τ_m,a_* and *τ_m,h_*, and translation rates *k_TL,a_* and *k_TL,h_*, whose values have been determined previously [32], and the threshold constants *K_a_* and *K_h_*, and Hill coefficients *n_a_* and *n_h_*, which can be estimated from bulk titrations (Fig. 1(d)). As a caveat, one has to consider that parameters measured in isolation do not necessarily match their apparent (effective) values in the coupled system [33] - for instance, we do not explicitly account for reactions such as the competition between *σ*^28^ and *σ*^70^ for the RNAP core enzyme [20, 34].

To illustrate the effect of system and control parameters on the dynamics of the free oscillator, we can consider the nullclines (*ṙ_a_* = 0, *ṙ_h_* = 0, with *ȧ* = 0, *ḣ* = 0, and assuming *δ* ≪ 1/*τ_m_*)

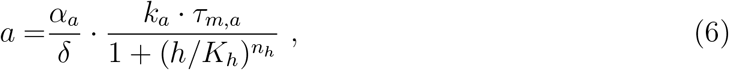

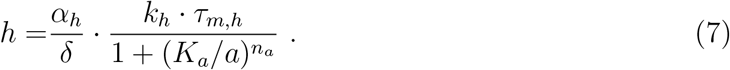

Hence, the system parameters describe the *shape* of the gene transfer functions, whereas the control parameters define their *scale*. As the stability of the fixed point at the intersection of the nullclines depends on the local shape of the nullclines (cf. the thorough linear stability analysis described in Ref. [15]), the control parameters can be used as bifurcation parameters to tune the qualitative behavior of the system, whereas the system parameters define the relative sizes of regions in parameter space corresponding to qualitatively different dynamics. For instance, increasing *n* will increase the parameter range that supports sustained oscillations. In the following, we assume that the system parameters are uniform for the activator and inhibitor, i.e., *α*:= *α_a_* = *α_h_*, *n*:= *n_a_* = *n_h_*, *k_TL_*:= *k_a_* = *k_h_*, *τ_m_*:= *τ_m,a_* = *τ_m,h_*, which is a standard approach to simplify the analysis while preserving the main qualitative features [12, 35].

### B. Operation of the free oscillator

To experimentally verify the predictions of the model, we tested the free oscillator for a wide range of dilution rates *δ* and transcription rates *α* (Fig. 2(a)). In good qualitative agreement with the model, we find regimes that display sustained, damped and strongly damped oscillations with varying periods. For the simulations, we used *α* as a global fitting parameter with a fixed ratio between samples. Reduction of *α* leads to a transition from sustained to damped oscillations, whereas *δ* mainly affects the period of the oscillations.

**FIG. 2.**
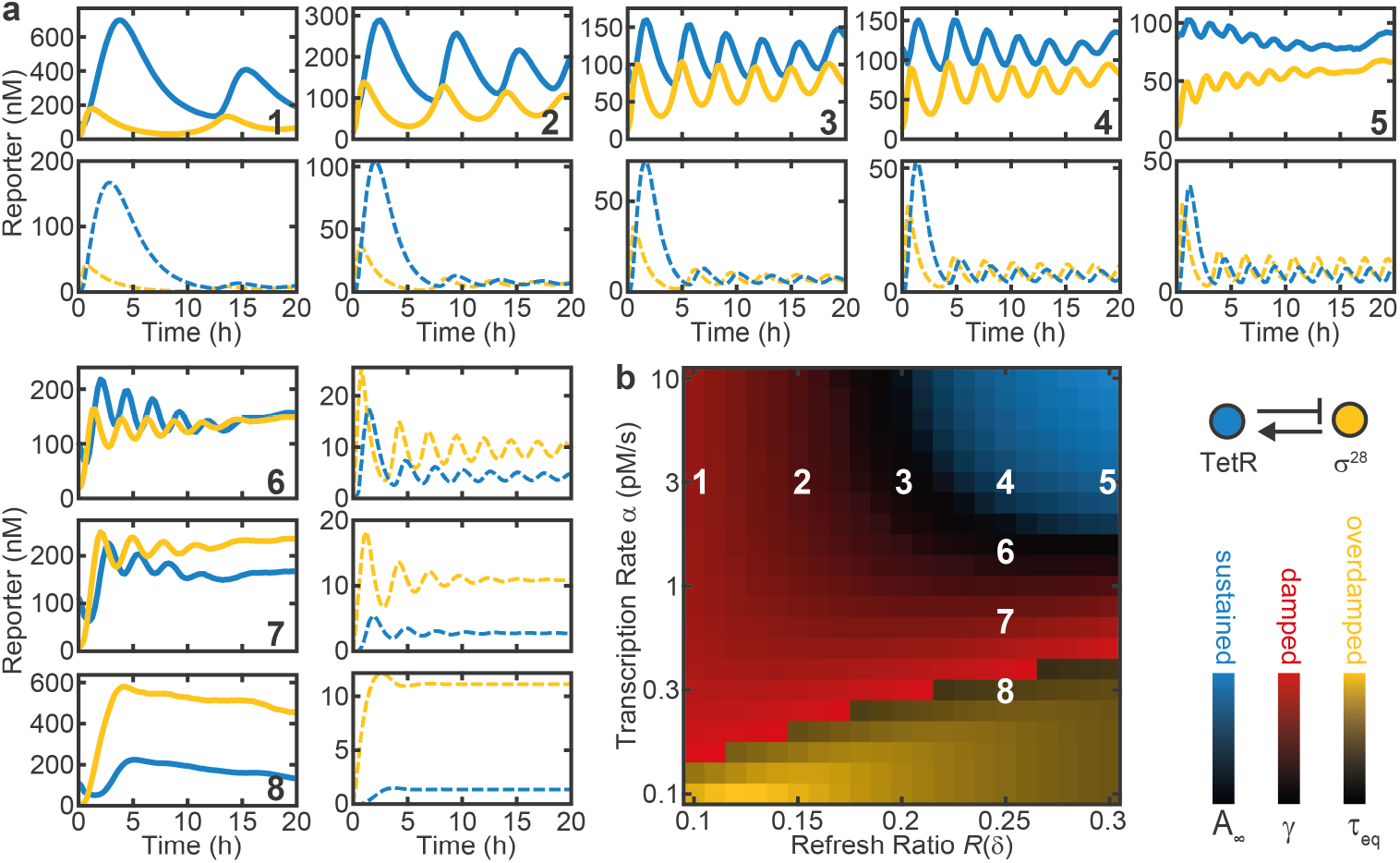
Dynamics and phase diagram of the free oscillator. (a) Experimental (full) and simulated (dashed) time traces of the free oscillator for different refresh ratios *R*(*δ*) and DNA concentrations as indicated in (b). (b) Simulated phase diagram of the free oscillator. Color overlay of different metrics as indicated in the legend reveals regions of sustained, damped and overdamped oscillations. *A_∞_*, γ, and *τ_eq_* are the normalized equilibrium amplitude, damping ratio, and equilibration time constant for trajectories with less than one detectable maximum, respectively. Experimental data was mapped onto the diagram using *α* as a fitting parameter proportional to the DNA concentration. System parameters: *n* = 3, *K_a_* = 20 nM, *K_h_* = 2 nM, *k_TL_* = 0.02 s^-1^, *τ_m_* = 12 min. DNA concentrations are 0.1–1 nM (0.3–3 pMs^-1^) for the circuit plasmids and 2 nM for reporter plasmids.

We also mapped the oscillator dynamics onto a simulated phase diagram, as shown in Fig. 2(b). To this end, we characterized the time traces of numerically simulated oscillations by their equilibrium amplitude *A_∞_* and mean damping ratio 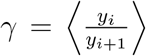 calculated from successive maxima *y_i_*. Strongly damped oscillations with less than two maxima were characterized by the exponential equilibration time *τ_eq_*. This illustrates that higher gene expression strength *α* and higher dilution rates *δ* favor sustained oscillations.

### C. Intrinsic timescale of the oscillator

We next investigated the dependence of the period *T* of the free oscillator on the model parameters using a simple form of sensitivity analysis (Fig. 3(a)). To this end, we tested the change of the period Δ*T* in response to a 30% change in each of the parameters individually [9]. In agreement with our naive expectation, the dilution rate *δ* is found to be the dominant control parameter determining the period of the oscillator *T*. The only other relevant parameters are the Hill coefficients *n_a_* and *n_h_*, and the mRNA lifetime *τ_m_*, which are fixed system parameters.

**FIG. 3.**
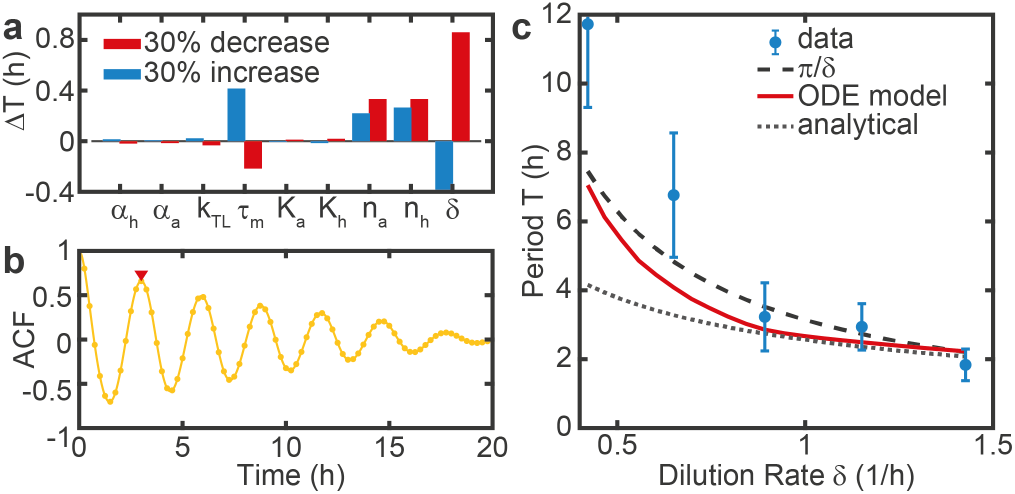
The period of the free oscillator is determined by the dilution rate. (a) A sensitivity analysis was performed by analyzing the change in the period Δ*T* in response to a 30% change in individual model parameters [9]. (b) Example of an auto-correlation function (ACF) that was used to estimate experimental periods for both reporters, corresponding to sample 4 in Fig. 2(a). The red triangle indicates the first maximum. (c) Period as a function of the dilution rate *T*(*δ*). The experimentally measured periods are compared to predictions by our ODE model, an analytical solution by [35], both with an mRNA lifetime of 12 min [32], and with the phenomenological *T*(*δ*) = *C_π_*/*δ* with *C* ≈ 1. While the predictions are in good agreement with the data at higher dilution rates, the measured periods are systematically higher at lower dilution rates. Error bars are standard deviations of 2-4 measurements, plus a systematic uncertainty that scales inversely with the number of maxima in the recorded time trace. System parameters are as in Fig. 2, with *α* = 3 pMs^-1^.

We then estimated the oscillator periods from experimental data using the first maximum of the auto-correlation function (ACF). Fig. 3(b) shows an example ACF corresponding to sample 4 in Fig. 2(a). The experimental data agrees well with the predictions from our ODE model for different *δ*, as shown in Fig. 3(c). The discrepancies at lower dilution rates are likely a result of the low number of complete cycles in the experimental time traces due to the long periods. This leads to an overestimation of the experimental periods, as the system initially has to approach the limit cycle. In contrast, an analytical solution (Eq. (24) in Ref. [35]), appears to more strongly underestimate the periods at lower dilution rate, probably as a result of the assumption that time traces are sinusoidal. Phenomenologically, we find that the ODE model predictions and the experimental data can be well approximated by the simple equation

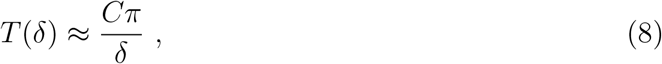

where *C* has a value close to 1. Importantly, this allows us to define the systems timescale based on the dilution rate *δ*.

### D. External forcing of the genetic oscillator

Next, we investigated whether we can force the oscillator to adapt to a certain period by externally supplying a periodic input signal. To this end, we replaced either the cell extract or the buffer supplied in every *k*-th dilution step with extract or buffer supplemented with TetR or aTc, to repress or activate the expression of *σ*^28^, respectively (Fig. 4). This generates a periodic input signal with a period *T_in_* = *k* · *t_int_* and amplitude *A_in_*, that rises instantly and decays exponentially with rate *δ*, as monitored with a fluorescent reference signal (mScarlet-I).

**FIG. 4.**
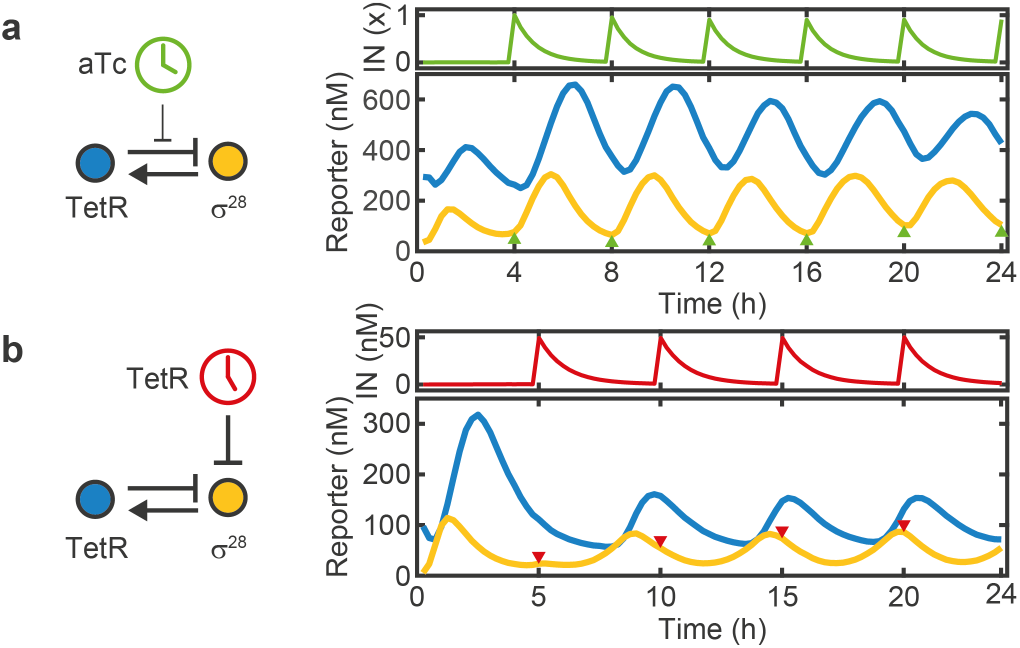
Dynamics of the forced oscillator. Here, we either added (a) 1× = 214 nM aTc every 4 h (λ = 1.43), or (b) 50 nM TetR every 5 h (λ = 1.48). Once added, the input signals are removed by the dilution with fresh reagent (without input), which results in an exponential decay of the input signal. The oscillator responds by increasing, or decreasing the production of *σ*^28^, respectively, and synchronizes to the input period.

The amplitude of the input signal must be chosen such that it triggers a sufficiently strong response by the oscillator, but is also diluted to levels well below the induction threshold sufficiently fast. For instance, an input signal with amplitude 1 will drop to (1 – *R*)^*k*^ = 0.01 after *k* = 16 dilution steps with a refresh ratio *R* = 0.25. Hence, in practice there is a minimum input period, typically ≈ 2 h, below which effective forcing becomes challenging due to the low attainable dynamic range of the input signal.

As shown in Fig. 4, the forced oscillators quickly adapt to *T_in_* within ≈ 1 – 2 cycles for both methods of external driving. While the amplitude is enhanced for positively forced oscillations, it decreases for negatively forced oscillations and the phase of the *σ*^28^ signal is shifted by ±*π*/2 relative to the input signal for positive and negative forcing, respectively. In both cases the system displays regular 1-cycle oscillations.

As described below, the forced oscillator system can exhibit more complex dynamics, which can be described with a single dimensionless bifurcation parameter

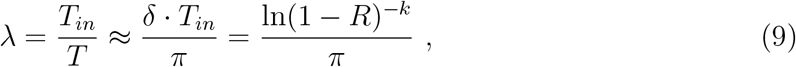

which is obtained as the ratio of input period and intrinsic period given in Eq. (5) and Eq. (8) (with *C* = 1), respectively. Note that ln(1 – *R*)^-*k*^ is the logarithm of the total dilution after one input period.

### E. Period doubling

In the following, we focus our analysis on the negatively forced oscillator that uses TetR as the periodically varying external input. When simulating the forced oscillator for different natural periods *T* and and input periods *T_in_*, we found that under certain conditions the resulting oscillations displayed a sequence of maxima with varying height that repeated every two or four maxima. Such a period doubling phenomenon commonly appears in non-linear systems of coupled or forced oscillators [29] and is a well studied route towards chaos [36]. Even though being investigated in great theoretical detail, period doubling has not been experimentally demonstrated in the context of synthetic biochemical oscillators so far [9]. Two of the experimental challenges in this context are that to record an *m*-cycle, the oscillator has to run reliably for *t* > *T_in_* · 2 · *m* = 32 h (for *T_in_* = 4 h and *m* = 4), and that for increasing *m* the bifurcation parameter λ has to be tuned with an exponentially increasing accuracy [37].

As shown in Fig. 5(a), we indeed find experimental evidence of period doubling in our system (here for λ = 1.20). After the typical large first maximum that occurs during the initial transient, the forced oscillator approaches a 4-cycle and stays there for two full revolutions. In the experiment, period doubling is more evident in the TetR dynamics than in the *σ*^28^ dynamics. After about 36 h the system appears to ‘drop back’ to a regular 1-cycle. As 36 h is close to the longest time span for which such a reactor was reportedly operated [28], this behavior is likely a sign of fatigue, which is also consistent with an observed drop of the refresh ratio towards the end of the recording.

**FIG. 5.**
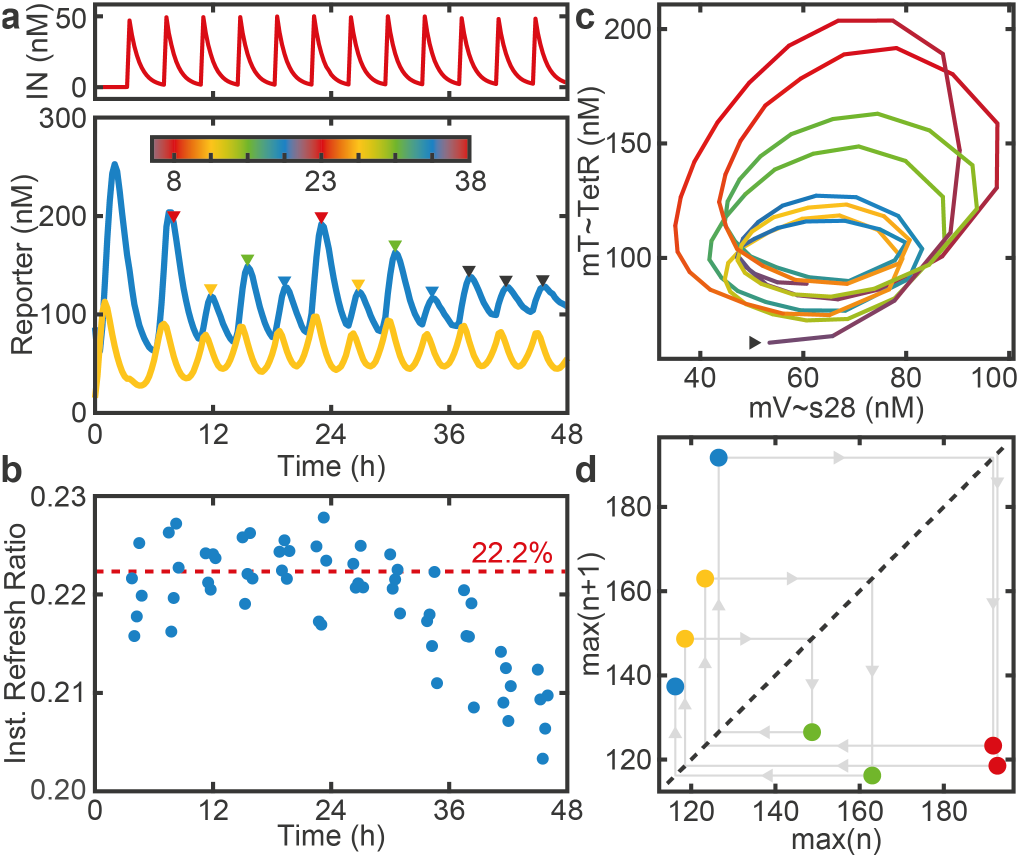
Period Doubling. (a) Time traces of a 4-cycle (λ = 1.20). (b) The instantaneous refresh ratio drops after 36 h due to fatigue. The dashed line indicates the average *R_t_* over the first 24 h. (c) Phase space trajectory. Color indicates time and is scaled to span 4 · *T_in_*, as indicated by the colorbar in (a). The arrow marks the starting point of the trajectory, which follows a counterclockwise path. (d) Maximum return map for the maxima indicated in (a). After the last point, the system transitions to a 1-cycle.

We computed an ‘instantaneous refresh ratio’ *R_t_* = 1 – *I_t+1_*/*I_t_* (Fig. 5(b)) using the reference time traces *I_t_* for all time points *t* where *I_t_* > 0.3 · *I_max_*. For *t* ≳ 36 h, *R_t_* drops by about 1%, leading to a decrease in δand a corresponding change in λ. Note that the instantaneous refresh ratio also slightly deviates from the nominal refresh ratio (here 20%) that was defined by calibration prior to the experiment. We hence use the more accurate instantaneous refresh ratio to calculate the control parameter *λ*. Similarly, a loss in activity of the supplied reagents would lead to a decrease in *α* over time, resulting in a stronger damping of the free oscillations.

Period doubling can further be visualized with a phase portrait (Fig. 5(c)), which highlights that the trajectories return to their starting point in phase space after completion of four revolutions. Finally, we can generate a maximum return map (Fig. 5(d)) by plotting the amplitude of each maximum against that of its predecessor. Again it can be observed that, within experimental variability, the system visits four distinct points in the map until it returns to its original location.

### F. Bifurcation diagram

In order to gain a more complete overview of the dynamical repertoire of our biochemical oscillator, we simulated a bifurcation diagram (Fig. 6(a)), for which we plotted the heights of the maxima against the parameter λ. Because successive bifurcations occur within exponentially decreasing intervals [37], we adjusted λ(*R*) as smoothly as possible. Following Eq. (9), we fixed the input period to *T_in_* = 4 h, as it can only be adjusted in increments of *t_int_* ≥ 15 min, and instead varied the natural period *T* by adjusting the refresh ratio *R*, which can be varied, in principle, continuously.

**FIG. 6.**
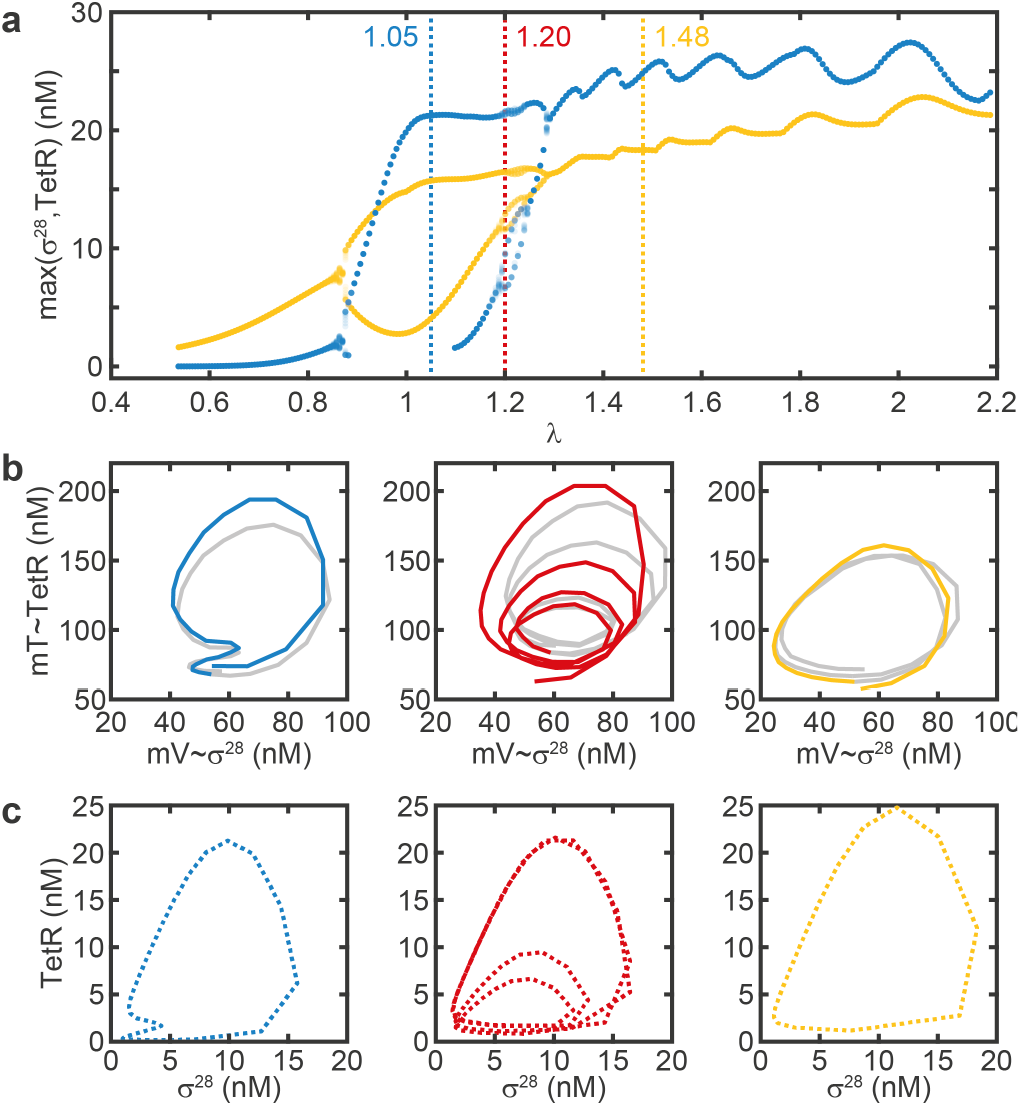
Bifurcation analysis for varying λ. (a) Simulated bifurcation diagram showing the maxima for *σ*^28^ (yellow) and TetR (blue). The values of λ that correspond to the experimental data in (b) and simulations in (c) are indicated by dashed vertical lines. (b) Experimental phase portraits exhibit a 2-, 4-, and 1-cycle. A second (third) *m*-cycle is grayed out for clarity. The 2-cycle is degenerate in the TetR-axis, i.e. the second maximum does not show in the time traces. Data for the 1-cycle and 4-cycle is the same as shown in Fig. 4b and Fig. 5. (c) Simulated phase portraits. Parameter values are as in Fig. 2, except *α* = 5 pMs^-1^, *n* = 4, *T_in_* = 4 h, *A_in_* = 50 nM.

The bifurcation analysis reveals that for low λ the system displays 1-cycles, then undergoes two period doublings to show a 4-cycle around λ = 1.2, followed by period ‘halvings’ and again 1-cycles at high values of λ. Qualitatively, the experimentally recorded phase space trajectories (Fig. 6(b)) match the corresponding simulated trajectories for the same λ values (Fig. 6(c)) remarkably well. One interesting feature is that for λ = 1.05 both experiment and simulation display a 2-cycle that is degenerate in the dynamics of TetR, i.e., the second maximum is not visible, but the period is doubled.

### G. Chaotic dynamics in the oscillator model

We were finally interested whether our system could, in principle, exhibit even more complex dynamics than a 4-cycle. We therefore simulated a two-dimensional bifurcation diagram, for which we varied both the Hill coefficient *n* and λ (Fig. 7(a)). The system dynamics can then be classified by means of the rotation number *m*, which equals the number of periods the system undergoes before returning to the starting point. For a chaotic trajectory, *m* = ∞, but for practical reasons we classify trajectories as chaotic if *m* > 32. As shown in Fig. 7(a), the oscillator model indeed permits chaotic solutions. Notably, chaotic regimes are interrupted by windows of mostly period 3, which is a commonly observed phenomenon [29]. The existence of *m* = 3-cycles actually implies the existence of chaotic trajectories [38], examples of which are shown in Fig. 7(b,c). Overall, this analysis reveals that higher order period doublings and chaotic behavior become increasingly prevalent for increasing non-linearity, corresponding to increasing Hill coefficients in the oscillator model.

**FIG. 7.**
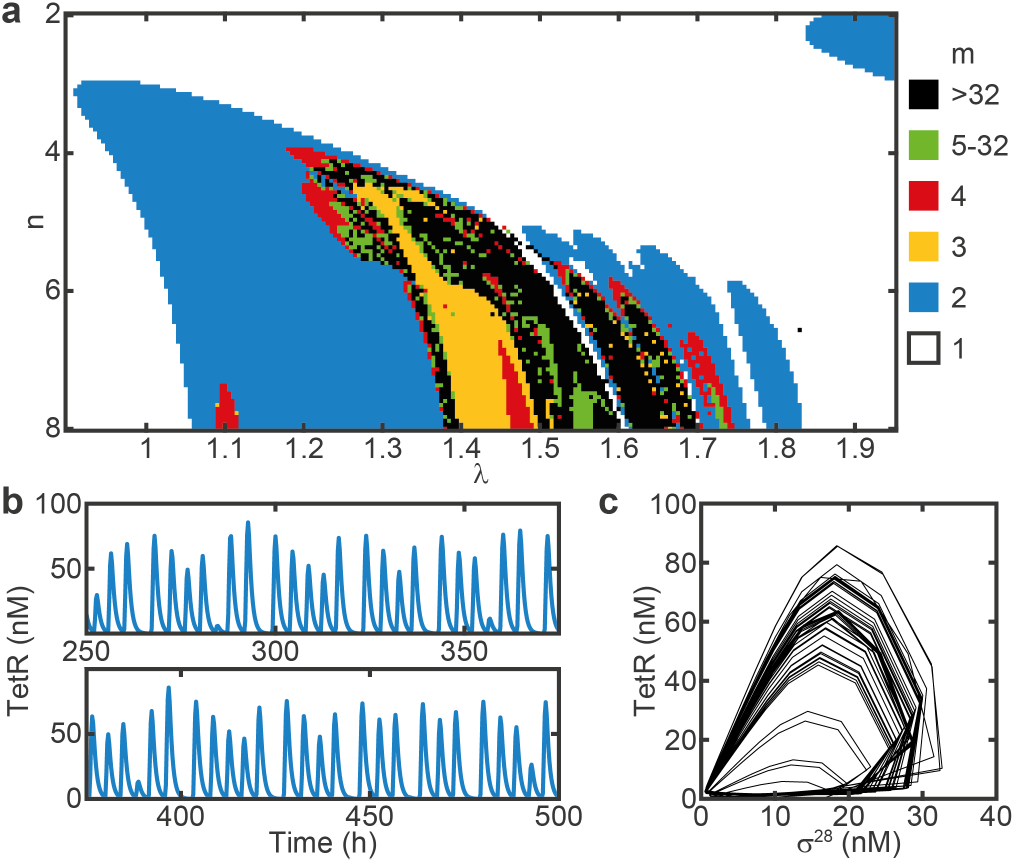
Simulations predict chaotic dynamics. (a) Two-dimensional bifurcation diagram showing the rotation number *m* for varying *n* and λ. With increasing *n*, the dynamics become increasingly more complex, over a broader range of λ. (b) Chaotic trajectory and (c) phase portrait of the corresponding strange attractor for *n* = 8, λ = 1.5542. Parameter values, except *n*, are the same as in Fig. 6.

## III. CONCLUSION

Inspired by the entrainment of biological circadian clocks by environmental zeitgeber cues, we have here experimentally investigated the response of a single-loop cell-free genetic oscillator to externally applied periodic perturbations. To this end, we utilized a microfluidic reactor system which allowed precisely controlled addition of components to the oscillator and dilution at regular intervals. The period of the free-running oscillator is dominated by the reactor’s dilution rate *δ*, which defines a timescale 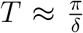. External forcing of the oscillator is achieved by periodically supplying transcription factors or inducers with an input period *T_in_*. The dynamic behavior of the forced oscillator is then determined by a single bifurcation parameter λ, which is defined as the ratio of the input period and the intrinsic timescale, i.e., λ = *T_in_*/*T*. Supported by simulations, we experimentally found non-trivial behavior such as 2- and 4-cycles, which is indicative of period doubling. Complex dynamics was observed for λ close to 1, whereas higher values of λ = 1.48 lead to stable entrainment of the genetic oscillator to the zeitgeber timescale.

To our knowledge, period doubling has not been found in experiments with biological circadian clocks so far, but has been repeatedly predicted by theoretical models. In a theoretical study by Kurosawa and Goldbeter, a tendency towards complex dynamics for λ ≈ 1 was found in a model of the *Neurospora* clock for strong forcing amplitudes [7]. However, this behavior was not found in a model of the *Drosophila* circadian clock by the same authors. The authors attribute this to differences in the forcing mechanisms, which in *Neurospora* alters expression rates, whereas in *Drosophila* alters degradation rates. Also, the authors argue that the complexity observed for λ ≈ 1 may explain why the free-running periods of many circadian rhythms differ substantially from a 24 h period. For instance, in humans, the free-running circadian period is 24: 11 h ± 0: 08 h (± SD), but in simpler organisms varies from 21.5 h (λ = 1.12) in *Neurospora* [39] to about 28 h (λ = 0.86) in *Phaseolus* [40].

Both from a biological point of view as well as for potential applications of synthetic biochemical clocks it might be desirable to actually prevent complex behavior resulting from coupled oscillator dynamics, which can be achieved in various ways. One possibility is to simply tune the free-running period away from the zeitgeber period, and thus avoid λ ≈ 1.

We further notice from our simulated bifurcation diagram Fig. 6(a) that as λ increases, the system initially undergoes two period doublings, but then does not continue to produce chaos, but follows with period halvings that eventually lead to stable 1-cycles. Similar period-halvings have been found in the study of ‘single-humped’ 1-dimensional maps [41], which are used as simple ecological models, when the recursion map was modified with a constant ‘immigration’ term that prevented the population from ever falling below a certain floor level. In the context of our biochemical oscillator, such a term would correspond to leaky/basal gene expression, potentially having a similar effect on the reversal of period-doubling transitions.

While in biological systems, biochemical rhythms are subject to evolutionary optimization, in synthetic biological systems robust behavior can be achieved by appropriately tuning system parameters such as the shape of the gene transfer functions [33]. Such engineering may be supported by *in silico* modeling, e.g., using evolutionary algorithms that intrinsically generate robust solutions [42], combined with high-throughput microfluidic reactors that enable comprehensive parameter screens [43].

In summary, we have shown that a synthetic cell-free gene circuit operated in a microfluidic reactor can be used to physically emulate the entrainment of a genetic oscillator with an external zeitgeber signal, which allowed the experimental realization of periodic doubling bifurcations, which had been previously only observed in numerical models of such systems. Apart from the fundamental interest in oscillatory biochemical systems, synthetic biochemical clocks may be of use in a wide range of applications that require intrinsic time measurements for the autonomous orchestration of downstream processes. In order to improve the accuracy of such oscillators and synchronize them to an external clock, coupling to a zeitgeber signal will be required. Our study demonstrates how the choice of system and control parameters can be used to tune the dynamics of such systems to become robust - or complex.

## IV. METHODS

### A. Microfluidic chip fabrication

The microfluidic reactor used in this study was fabricated with multilayer photo- and soft-lithography methods as detailed in [44]. The structures on the control layer master were patterned from 40 μm SU8-3050 (micro resist technology). To reduce the minimal refresh ratio per feed *R*_0_ (≥ 0.3% vs. ≥ 2%), we increased the volume of the ring reactors ≈ 10-fold using a 2-layer design (50 μm SU8-3050 and 20 μm AZ 40XT (MicroChemicals)) for the flow layer master, similar to [16]. The structures on the flow layer master were enlarged by 1.8% to correct for shrinking of the PDMS relative to the control layer. All masters were treated with trichloro(1H,1H,2H,2H-perfluorooctyl)silane (Merck, #448931-10G) in a weak vacuum for at least 2 h.

The PDMS device was fabricated by first casting an appropriate amount of PMDS, mixed at a 5:1 ratio with crosslinker, onto the flow layer master. The control layer was prepared by spin-coating PDMS (20:1) onto the master to a height of 50 μm. After relaxation for 45 min, the molds were baked at 80 °C for 20 min, or 25 min, respectively. The flow layer was removed from the master, trimmed, and aligned on the control layer using a stereomicroscope. After thermal bonding at 80 °C for 90 min, devices were trimmed and holes for control and flow lines were punched using catheter punches (Syneo, #CR0320245N21R4). Finally, devices were cleaned with Scotch tape and plasma-bonded onto a clean glass slide.

### B. Cell-free gene expression reactions

Homemade *E. coli* cell extract was prepared from Rosetta 2(DE3) by sonication based on standard protocols [45, 46]. The final reaction contained 50 mM Hepes pH 8, 1.5 mM ATP (Roth, #HN35.3) and GTP (Roth, #K056.4), 0.9 mM CTP (Roth, #K057.4) and UTP (Roth, #K055.3), 0.2 mg mL^-1^ tRNA (Merck, #10109541001), 26 mM coenzyme A (Merck, #C3144-10MG), 0.33 mM NAD^+^ (Merck, #481911), 0.75 mM cAMP (Merck, #A9501-1G), 68 μM folinic acid (Merck, #47612-250MG), 1 mM spermidine (Merck, #S2626-1G), and 30 mM 3-PGA (Merck, #P8877-1G), as an energy source. The final concentrations of screened components were [46] 4 mM Mg-glutamate (Merck, #49605-250G), 60 mM K-glutamate (Merck, #49601-100G), 1.5 mM of each amino acid except leucine (Biozym, #BR1401801), 1.25 mM leucine, 2.5 % (w/v) PEG-8000 (Merck, #89510-250G-F), and 0 mM DTT. A final concentration of 2 mM TCEP (Roth, #HN95.1) was added to the buffer solution immediately prior to the experiment to allow storage of buffer reservoirs at ambient temperature [47].

DNA templates were assembled from various sources (Biobricks, IDT gBlocks, *σ*^28^ was PCR amplified directly from the genome of *E. coli* strain MG1655) using a standardized Golden Gate Assembly scheme [31] and cloned into DH5*α* or DH5*α*Z1, when using TetR repressible promoters. Plasmids for expression were prepared using a Midiprep kit (Qiagen, #27104) and concentrations were estimated by UV-Vis spectroscopy. DNA sequences are available upon request.

Added proteins (TetR, mTurquoise2, mVenus, mScarlet-I) were purified using standard His-tag Ni-NTA affinity chromatography. Briefly, the gene of interest was cloned into a 6xHis-pSB1A3-pT7 expression vector and expressed in BL21star(DE3). A 500 mL culture was harvested, lysed via sonication and purified using HisTrap HP columns (GE, #17-5247-1). The fractions were analyzed with SDS-PAGE, pooled and concentrations were estimated by UV-Vis spectroscopy.

Reactions were prepared in separate tubes for extract and buffer and combined on chip in a 1:1 ratio, usually consuming 45 μL of each reagent solution per 24 hours. Additional proteins were added to the extract, whereas DNA or other components were added to the buffer.

### C. Experimental setup and operation

The experimental setup used to control the microfluidic device and image acquisition was custom built around an Olympus IX81 inverted epifluorescence microscope equipped with a motorized stage, fluorescence light engine (Lumencor SOLA SE II 365), camera (Andor iXon3 DU888), and filters (CFP: 438-25/458/483-32, YFP: 500-20/515/535-30, RFP: 559-34/588/609-34, GFP: 472-30/495/520-34). The reaction temperature was kept at 29 °C using a cage incubator (Okolab).

Extract reservoirs were kept at 4 °C with a custom built cooling unit fit for two 1.5 mL Eppendorf tubes. Flow line pressures were regulated to 300 mbar using a pressure controller (Elveflow OB1). Control lines were operated with 1.5-2.5 bar using a custom built valve controller based on 24 solenoid valves (Festo, #MH1) and an Arduino Mega. Feed, mix and acquisition cycles during time lapse were automated with a custom LabVIEW program, which allowed execution of arbitrary input programs. All chips were calibrated to determine Ro for each reactor with 25 μM fluorescein in PBS prior to the experiment.

### D. Data analysis

Microscope images were analyzed with custom matlab scripts. First, an ROI and background ROI were manually selected for each ring and the background subtracted average intensity *I* – *B* was normalized against the corresponding measurement from the *R*_0_ calibration *I*_0_ – *B*_0_. Using similarly generated 1-point reference measurements *I_ref_* – *B_ref_* (1 μM of reporter protein in cell extract) and *I*_0,*ref*_ – *B*_0,*ref*_ (25 μM fluorescein in PBS), we obtain a reporter concentration c, which is comparable across experiments

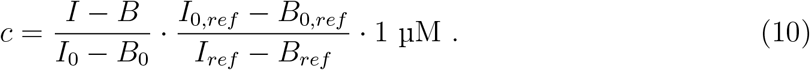

Maxima were detected using *findpeaks* and auto-correlation was computed using *xcorr*. The rotation number m was determined by computing the auto-correlation of the maxima.

### E. Modeling

Simulations were performed in matlab using *ode23s* to solve the ODE model during the interval of one dilution cycle *t_int_* = 15 min. After each cycle, the initial conditions of the consecutive cycle *c_i+1_* were set with the final concentrations of the previous cycle *c_i_* as *c_i+1_* = *c_i_* · (1 – *R*) + *c_in_*, where *R* is the refresh ratio and the input period *c_in_* = *A_in_*, if *i* mod *k* = 0, and *c_in_* = 0, otherwise. To approximate the experimental procedure, we sample the solutions at the final time point of each interval.

## ACKNOWLEDGEMENTS

We would like to thank Zoe Swank, Sebastian Maerkl, and Nadanai Laohakunakorn for sharing their valuable time and knowledge on setup, device manufacturing, and operation of the microfluidic reactor. Further, we wish to thank Elisabeth Falgenhauer and Aurore Dupin for their advice on cell extract production, purification of TetR, mTurquoise2, and mVenus, and providing several gene fragments, and Matthaeus Schwarz-Schilling for early discussions on synthetic genetic oscillators. This project was funded by the European Research Council (project AEDNA, grant agreement no. 694410).

